# Hierarchical control of cardiomyocyte maturation and ischaemia sensitivity by metabolic culture conditions

**DOI:** 10.64898/2026.03.12.711459

**Authors:** Yuanzhao Cao, Chris Siu Yeung Chow, Sumedha Negi, Woo Jun Shim, Sophie Shen, Chen Fang, Nathan J. Palpant

## Abstract

Ischaemic heart disease remains the leading cause of mortality worldwide, yet no therapies prevent cardiomyocyte death during acute ischaemia–reperfusion injury (IRI). Human induced pluripotent stem cell–derived cardiomyocytes (hiPSC-CMs) provide a platform for modelling cardiac injury, but their immature phenotype limits the physiological fidelity of *in vitro* models. Here, we systematically evaluated how experimental variables used during preparation of hiPSC-CM endpoint assays influence cardiomyocyte maturation and susceptibility to IRI. Integrating literature mining, molecular profiling, statistical genetics, and functional assays, we examined the effects of replating conditions, backbone media, metabolic substrates, and signalling modulators. We define the relationship between culture conditions and metabolic supplements in determining contractile maturation and sensitivity to IRI. Notably, we show that metabolic composition of the backbone medium establishes the baseline maturation state and determines responsiveness to additional maturation cues. These findings identify metabolic environment as a dominant regulator of injury susceptibility and provide a framework for improving the physiological fidelity of hiPSC-CM models of cardiac ischaemia.

## Introduction

Ischaemic heart disease (IHD) remains the leading cause of mortality worldwide. Acute myocardial infarction (MI), caused by coronary artery occlusion and subsequent ischaemia–reperfusion injury (IRI), results in extensive cardiomyocyte death and adverse cardiac remodelling. Despite substantial advances in reperfusion therapies, no clinically approved drugs effectively prevent cardiomyocyte loss during the acute phase of ischaemic injury^1,2^. This persistent translational gap highlights the need for physiologically relevant human models that accurately capture the determinants of cardiomyocyte vulnerability to ischaemia.

Human induced pluripotent stem cell–derived cardiomyocytes (hiPSC-CMs) provide a powerful platform for modelling human cardiac disease and enabling mechanistic and preclinical studies in a patient-specific context^3,4^. However, hiPSC-CMs resemble fetal rather than adult cardiomyocytes in several key aspects, including myofilament composition, contractile properties, calcium handling, and metabolic phenotype^5^. Adult cardiomyocytes primarily generate ATP through mitochondrial oxidative phosphorylation, whereas immature hiPSC-CMs exhibit reduced expression of tricarboxylic acid (TCA) cycle and fatty acid β-oxidation pathways and rely predominantly on glycolytic metabolism^6^. This metabolic divergence has important implications for modelling ischaemic injury, as glycolysis-dominant cells are generally more tolerant of hypoxic stress, while oxidative adult cardiomyocytes are more susceptible to energetic failure and cell death during ischaemia–reperfusion^7-9^. Consequently, immature hiPSC-CMs may underestimate injury severity and limit the physiological relevance of *in vitro* IRI models.

Substantial effort has therefore focused on improving the maturation of hiPSC-CMs for modelling cardiovascular disease^5,10,11^. Diverse strategies have been developed to promote maturation, including modulation of signalling pathways^12-14^, hormonal stimulation^15-17^, metabolic substrate manipulation^6,18,19^, and application of physiological cues^20-22^. While these approaches have advanced structural and functional maturation, the extent to which defined maturation conditions influence susceptibility to IRI remains incompletely understood. Here, we focus on the relationship between metabolic maturation and injury responsiveness in hiPSC-derived cardiomyocytes.

In practice, modelling ischaemic injury requires differentiated cardiomyocytes to be replated into endpoint assays, where experimental parameters such as cell replating density, extracellular matrix, backbone media, and metabolic supplementation can substantially reshape cardiomyocyte state. These stepwise protocol choices can alter cardiomyocyte state, including molecular programs associated with physiological maturation, metabolic phenotype, and susceptibility to hypoxic stress. In this study, we systematically evaluate these experimental variables across the workflow used to prepare hiPSC-CMs for ischaemia modelling. Specifically, we test how replating conditions, culture substrates, backbone media, and metabolic cues influence cardiomyocyte maturation and determine whether these states align with those that confer sensitivity to ischaemic injury. By perturbing metabolic substrates and media composition, we further examine context-dependent and hierarchical relationships between protocol variables that govern their utility for functional assays and disease modelling.

## Methods

### Generation and maintenance of human iPSC lines

All human pluripotent stem cell studies were carried out in accordance with consent from The University of Queensland’s Institutional Human Research Ethics Committee (UQ HREC approval #2015001434). Cardiomyocytes generated in this study were derived from the WTC-11 hiPSC line (Gladstone Institute of Cardiovascular Disease, UCSF)^23,24^. All WTC cells were maintained as previously described with slight adaptations^25,26^. Briefly, iPSC cells were maintained in mTeSR Plus medium with supplement (Stem Cell Technologies, Cat.#100-0276) at 37°C with 5% CO_2_. Cells were cultured on Vitronectin XF (Stem Cell Technologies, Cat.#07180) coated plates (Nunc, Cat.#150318). A complete reagent and resource Table is provided in Supplementary Information.

### CMPortal literature analysis of protocol variables

CMPortal^27^ (website link: palpantlab.com/cmportal) was developed as a public website source for hiPSC-CM protocol analysis. CMPortal corpus of 322 cardiomyocyte differentiation publications was analysed. Protocol variables, as previously^27^ defined, were sorted into five functional groups based on biological role: media, coating matrix and materials, molecules and growth factors, metabolic substrates and components, and protocol length and seeding conditions. Protocol variables that were temporal (features explicitly specifying day-by-day timing) were excluded from functional-group summaries. Associations with outcomes and purposes were determined bidirectionally as reported, were further filtered to retain only those strongly significant at *p* <0.01. Two frameworks were considered: (i) maturity outcomes (across structural, functional, electrophysiological, and molecular domains) and (ii) disease modelling applications spanning six pathological categories including genetic disorders (*n* =18), arrhythmic conditions (*n* =15), ischaemic infarction (*n* =9), dilated cardiomyopathy (*n* =6), hypertrophic cardiomyopathy (*n* =6), and fibrotic diseases (*n* =4). Enrichment within functional groups was compared between the two frameworks.

### CMPortal protocol analysis

#### Enrichment of protocol variables

The curated set of protocol variables spanning all five functional categories was tested for associations with the best contractile force and ischaemic modelling. Features were ranked by the average confidence, derived from entropy– and Gini index–based measures of association strength. The top 10 features were retained for each analysis. Confidence values were -log10 transformed, and significance was assessed independently for entropy and Gini metrics. **Pie charts of metabolic and ischaemic modelling**. The 322 papers were analysed specifically. First, publications with metabolic strategy (*n* = 29). Second, publications with *in vitro* ischaemic modelling (*n* = 9), including ischaemic and MI modelling. Third, specific metabolic techniques and media types were the focus of the ischaemic modelling publications (*n* = 9). **Metabolic component ranking across maturity aspects**. Six metabolic components (T3, IGF-1, palmitic acid, fatty acid, fatty acids and lipids, galactose) were evaluated across high-maturity endpoints. Within each endpoint, features were ranked by average confidence, then inverted to yield inverse rank scores (higher means stronger association). Significance was assessed by entropy and Gini metrics independently. Four categories were defined: None, Entropy-only, Gini-only, and Entropy & Gini. **Comparative media analysis**. The maturation efficacy of RPMI-1640 + B27/insulin (RPMI+B27) versus DMEM no glucose + fatty acids (DMEM+FA) was evaluated combined with various metabolic components and matrix coatings. In total, 25 target features were selected for analysis, comprising three hiPSC-CM matrix coating types (Matrigel *n* =163, Geltrex *n* =33, Vitronectin *n* =10), seventeen individual metabolic components (T3, fatty acids, palmitic acid, creatine, taurine, dexamethasone, L-carnitine, amino acids, galactose, insulin-transferrin-selenium, vitamin B12, biotin, ascorbic acid, Albumax, B27, KOSR, IGF-1), and 5 metabolic component categories (hormonal stimulation, sugars/carbohydrates, amino acids/derivatives, signalling pathway regulators, kinase inhibitors). Two media-specific datasets were constructed using CMPortal filtering. For each base medium (RPMI+B27 and DMEM+FA), protocols were first aggregated, and their reported z-score standardised across maturation endpoints. For each base media subset, the best-reported value for every indicator was selected directly from the standardised data. This is directional, where indicators with higher values reflect maturity (e.g., sarcomere length, contractile force, calcium flux amplitude, conduction velocity, protein expression ratios) and indicators with lower values reflect maturity (e.g., beat rate, calcium timing, resting membrane potential).

### Differentiation of iPSC-derived cardiomyocytes

Cells were differentiated according to a modified monolayer platform protocol previously described^25,26^. On day –1 of differentiation, hiPSCs were dissociated using Versene (Thermo Fisher Scientific, Cat.#15040066), plated onto Vitronectin XF-coated flasks (Nunc, Cat.#156367) at a density of 1.8 × 10^5^ cells/cm^2^, and cultured overnight in mTeSR Plus medium supplemented with 10 µM Y-27362 ROCK inhibitor (Stem Cell Technologies, Cat.#72308). Differentiation was induced on day 0 by first washing cells with PBS, then changing the culture medium to RPMI-1640 medium (Thermo Fisher Scientific, Cat.#11875093) containing 3 μM CHIR-99021 (Stem Cell Technologies, Cat.#72054), 500 μg/mL bovine serum albumin (BSA, Sigma-Aldrich, Cat.#A9418), and 213 μg/mL ascorbic acid (AA, Sigma-Aldrich, Cat.#A8960). After 3 days of culture, the medium was replaced with RPMI-1640 containing 5 μM XAV-939 (Stem Cell Technologies, Cat.#72674), 500 μg/mL BSA, and 213 μg/mL AA. On day 5, the medium was exchanged for RPMI-1640 containing 500 μg/mL BSA, and 213 μg/mL AA without any additional supplements. From day 7 and onward, the cultures were fed every other day with RPMI-1640 containing 2% B27 supplement plus insulin (Life Technologies Australia, Cat.#17504001). Spontaneous beating was typically observed between days 9 and 11 of differentiation.

### Flow cytometry

At the time of harvesting and replating on day 15 of differentiation, a subset of cells (approximately 1 × 10^6^) was set aside for flow cytometry analysis of cardiomyocyte purity of differentiated cell populations. Cells were fixed with 4% paraformaldehyde (Sigma-Aldrich, Cat.#158127-5G), permeabilised in 0.75% saponin (Sigma-Aldrich, Cat.#S7900), and labelled with Phycoerythrin (PE)-conjugated sarcomeric α-actinin (SA) antibody (Miltenyi Biotec Australia Pty, Cat.#130-123-773) or Cardiac Troponin T (Thermo Fisher Scientific, Cat.#MA5-12960) or PE-conjugated anti-human isotype (IgG) control (Miltenyi Biotec Australia Pty, Cat.#130-119-964). Stained samples were analysed using a FACS CANTO II (Becton Dickinson) system with FACSDiva software (BD Biosciences). Data analysis was performed using FlowJo software version 10.6.2, and cardiac populations were determined with population gating from corresponding isotype controls. For all experiments, only cell preparations with greater than 80% sarcomeric α-actinin-positive or Cardiac Troponin T-positive CMs were used.

### Replate of CMs and maturation induction

Differentiated CMs were replated on day 15 of differentiation for *in vitro* hPSC-CMs maturation protocols. Cells were dissociated using 0.5% trypsin, stopped with Stop Buffer (1:1 FBS in RPMI-1640), filtered with a 100 μm strainer, replated at a density of 1.58 × 10^5^ cells/cm^2^ in Vitronectin XF-coated experimental plates with replating medium (RPMI-1640, 2% B27 supplement plus insulin, 5% FBS, and 10 µM Y-27362) and cultured overnight. FBS and ROCK inhibitor-containing medium was replaced the following day (day 0) with RPMI+B27 medium (RPMI-1640 plus B27/insulin). From the replating day 0 to day 7 post-replate, different backbone media were used to maintain CMs, including RPMI+B27 and DMEM+FA. Different modulators and maturation factors were supplemented into the backbone media for the end assays. During the maintenance stage, media were refreshed every other day.

### CAGE-Sequencing

The CAGE-sequencing protocol was described previously^28^. Library Preparation: Library preparation was performed by the Genome Innovation Hub (GIH) and UQ Sequencing Facility at the Institute for Molecular Bioscience using the DNAFORM commercial CAGE preparation user guide with the addition of a qPCR quantification step to normalised input for pooling before second strand synthesis Sequencing: Sequencing with 5% PhiX spike-in was performed on an Illumina NextSeq500 using the following configurations: Read1 – 76bp, Index1 – 6bp. Loading concentrations were 1.6pM for the cardiomyocyte samples, using a modified denaturation and dilution protocol for low-concentration libraries. TSS Analysis: CAGE-seq data was analysed using the CAGEr pipeline^29^. Differential Expression Analysis: Promoters and genes were assessed for differential expression using DESeq2, as described^30^. Gene Ontology Analysis: Gene ontology (GO) analysis was performed using DAVID, with significance threshold set at FDR < 0.05, with only level 5 biological process, molecular function, and cellular components GO terms selected^31^.

### Single-cell disease relevance score (scDRS) analysis

Disease gene sets consisting of the top 1000 associated genes and their Z scores to the relevant summary statistics were generated using MAGMA^32^. We converted the hiPSC differentiation scRNA-sequencing dataset^25^ to Anndata format for subsequent processing with the command line interface scDRS pipeline^33^ (https://github.com/martinjzhang/scDRS). Individual cell-level disease risk scores were computed (**Figure 1a, left**), followed by group-level associations based on time point in the single-cell dataset (**Figure 1a, right**).

**Fig. 1.**
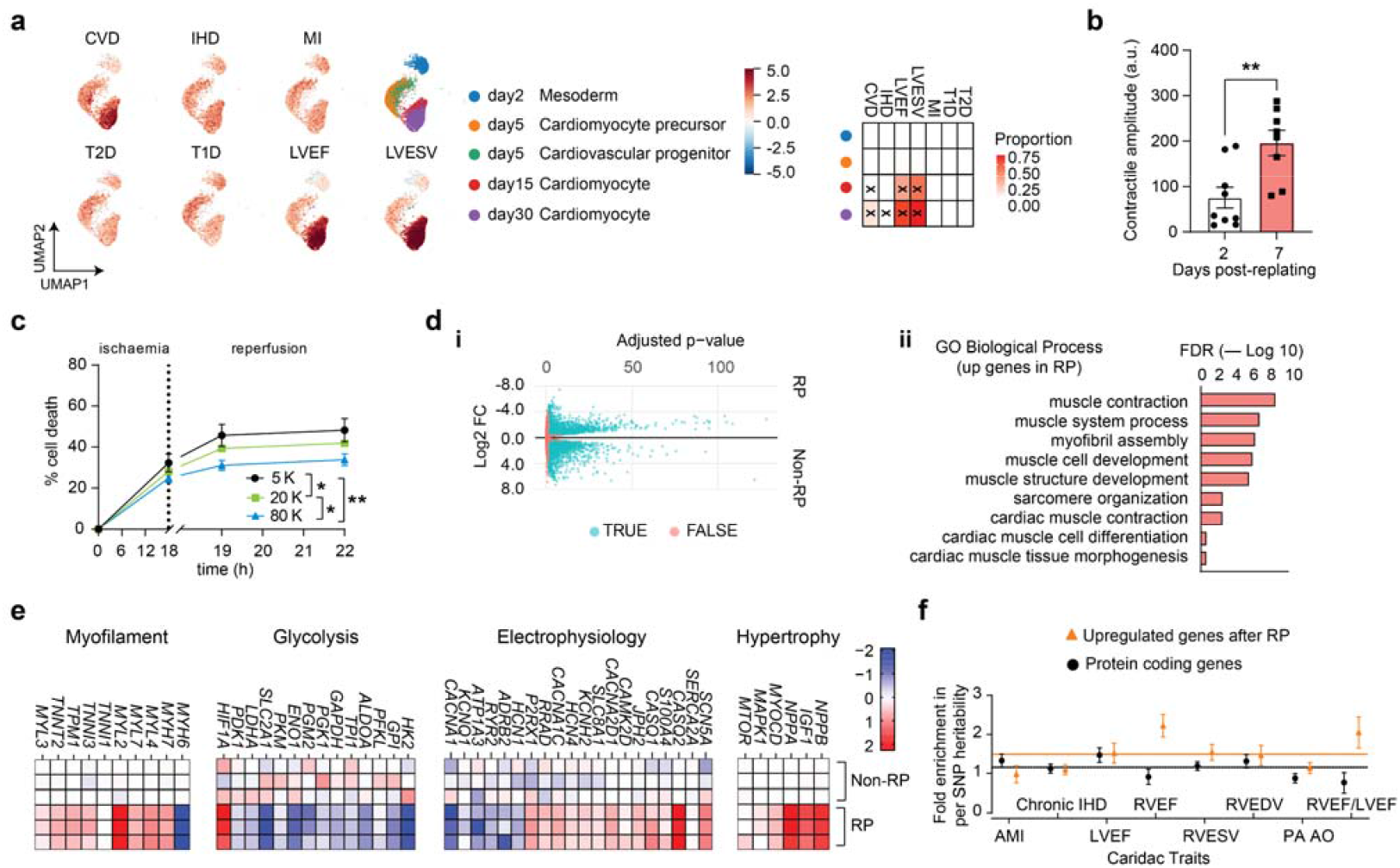
Comparative analysis of cell replating influencing iPSC-CM maturation. **(a)** scDRS analysis to compare molecular profiles of hiPSCs differentiated into the cardiac lineage at different time points with cardiovascular structural and functional traits, cardiovascular disease traits and diabetes-associated traits. x indicates significant scDRS enrichment between the cell type and the disease/functional traits. **(b)** MUSCLEMOTION analysis of 2D-based CM contractility amplitude under the condition of replating. **(c)** Analysis of CMs susceptibility to cell death under ischaemia-reperfusion stress evaluating the effect of three different CM replating densities. **(d)** CAGE-sequencing analysis of upregulated genes in replated CMs (RP) compared to non-replated CMs (Non-RP) shows a volcano analysis **(di)** and GO analysis **(dii)** of DE genes in RP CMs compared to Non-RP CMs. **(e)** Heat maps of gene programs associated with cardiomyocyte maturation between RP and Non-RP conditions. **(f)** Fold enrichment in per-SNP heritability (SNP-based heritability divided by the number of SNPs in the annotation relative to the whole genome) for up-regulated genomic regions after RP versus all protein-coding genomic regions. Fold enrichment was estimated by SBayesS, with the error bar being the posterior standard error. The analysed traits are human cardiac structure and ischaemia traits. T2D, type 2 diabetes mellitus; T1D, type 1 diabetes mellitus; MI, myocardial infarction; LVESV, left ventricular end-systolic volume; LVEF, left ventricular ejection fraction; IHD, ischaemic heart disease; CVD, cardiovascular disease; AMI, acute myocardial infarction; RVEF, right ventricular ejection fraction; RVESV, right ventricular end-systolic volume; RVEDV, right ventricular end-diastolic volume; PA AO, pulmonary artery and aorta diameter ratio; RVEF/LVEF ratio, inverse-normalised right ventricular ejection fraction to left ventricular ejection fraction ratio. Unless otherwise specified, *n* = 3; 3 biological replicates, each with 3 technical replicates. Data are presented as mean ± SEM. Statistical analysis by two-tailed student’s *t*-test **(b)**, or one-way ANOVA **(c)**. \**p*<0.05, \*\**p*<0.01, \*\*\**p*<0.001, \*\*\*\**p*<0.0001.

### Heritability analysis

SNP-based heritability was performed using SBayesRC^34^. Genes were mapped to ∼7M hapmap3 SNPs with a window of ±50kb to capture SNPs^35^ in the locus, including linkage disequilibrium regions and trans/cis regulatory elements, which the SBayesRC R pipeline was applied (https://github.com/zhilizheng/SBayesRC) with SNP imputation. Proportion of phenotypic variance in the traits explained by all protein-coding gene SNPs is estimated and compared to the proportion of variance explained by SNPs in genes upregulated after replating in RPMI+B27 backbone media and in genes upregulated after replating in DMEM+FA backbone media. Both scDRS and heritability analysis used UK Biobank GWAS summary statistics from either the Neale Lab (https://www.nealelab.is/uk-biobank) or summary statistics post-analysis UK Biobank cardiac MRI data^36,37^.

### Processing bulk RNA-sequencing data for gene quantification

hiPSC-CMs were cultured under backbone or perturbation conditions. Cells were subsequently harvested for RNA extraction and genome-wide RNA sequencing. RNA-sequencing samples with 3 biological replicates for each group were aligned to hg38 using STAR aligner^38^ with the default option, yielding more than 294M uniquely mapped reads in total with an average of 92.84% mapping rate. Gene-level read counts were quantified using htseq-count^39^ using the NCBI hg38 RefSeq gene annotation. Genes with count per million (CPM) of at least 1 in at least 2 samples were kept for further analysis. Read counts were normalised using the ‘calcNormFactors’ function in the EdgeR package, with TMM method^40^. Negative binomial generalised linear models were fit. Differentially expressed (DE) genes were identified by comparing all possible pair-wise combinations of the conditions (false discovery rate < 0.001).

### MUSCLEMOTION contractility analysis

Brightfield videos (AVI) of cardiomyocyte contractions were acquired at 60 frames per second and analysed with MUSCLEMOTION ImageJ plugin^41^.

### Contractile performance and CardioExcyte 96 recordings

Differentiated cardiomyocytes were replated on day 15 of differentiation for physiological maturation or culture conditions. 50,000 viable cells per well were plated on Vitronectin XF-coated 96-well CardioExcyte NSP-96 plates (Nanion Technologies GmbH, Cat.#201001). Contractile parameters of beating hiPSC-CM monolayers were recorded using the CardioExcyte 96 system (Nanion Technologies GmbH) for 7 continuous days after replating. Contractility recordings performed with CardioExcyte 96 were obtained with 1 ms time resolution and 1 kHz sampling rate. All experiments were conducted under physiological conditions using an incubation chamber integrated in the system (37 °C, 5% CO_2_, 80% humidity). The CardioExcyte NSP-96 plates utilise two gold electrodes on a rigid surface in each well to study physiological changes on contractility via continuous impedance monitoring of CMs exposed to maturation or stress conditions. The parameters for detection were amplitude, beat rate, and upstroke velocity.

### *In vitro* IRI model with hiPSC-CMs

Differentiated cardiomyocytes were replated on day 15 of differentiation for experimental conditions, followed by *in vitro* ischaemia assays. 40,000 viable cells per well were plated on Vitronectin XF-coated 96-well plates. After replating, the cells were recovered overnight in replating medium. The following day, the medium was replaced with the indicated maturation media conditions or factors, as specified in the experimental design. To prepare media for ischaemia/reperfusion injury, 10x HBSS without sodium bicarbonate (Sigma-Aldrich, Cat.# H6136) was diluted to 1x concentration in sterile tissue culture-grade water. Solutions were buffered with 12 mM HEPES (for pH 7.4 media, Sigma-Aldrich) and the pH adjusted accordingly with 1 M NaOH. The medium was sterile filtered with 0.22 μm syringe filters (Millipore). Unless otherwise noted, the replated cells were treated overnight (18 h) in HBSS under either normoxic (∼18.5% O_2_; 5% CO_2_) or hypoxic (0.5% O_2_; 5% CO_2_) culture conditions. For reperfusion experiments, the medium was refreshed with HBSS pH 7.4 after overnight incubation and cultured for 1 h and 4 h under normoxic conditions. To assess cell death, the supernatant was collected, and LDH levels were measured using a cytotoxicity detection kit (Roche, Cat.#11644793001). For all cell culture experiments, percent cell death was calculated using low and high controls. For low control, cardiomyocytes were cultured overnight in standard culture medium (RPMI + B27/insulin). For high control, cells were cultured in RPMI plus B27/insulin supplements containing 1% Triton X-100 (Sigma-Aldrich, Cat.#X100).

### Statistics and reproducibility

CMs on day 15 were used from 3 independent hiPSC differentiations. A minimum of three biological replicates was used for each experimental group. Statistical analysis was performed using Prism 9 version 9.3.1 (GraphPad Software). To compare two normally distributed groups, a two-tailed Student’s *t*-test was performed. For three or more groups and the assessment of one parameter, a One-Way ANOVA statistical test was used. With multiple parameters and three or more groups, a Two-Way ANOVA was used. Significance was determined as \**p* < 0.05, \*\**p* < 0.01, \*\*\**p* < 0.001 and \*\*\*\**p* < 0.0001. Data are presented as mean ± SEM.

## Data and code availability

Bulk RNA-seq has been deposited in the Gene Expression Omnibus database (GSE298573), the link is https://www.ncbi.nlm.nih.gov/geo/query/acc.cgi?acc=GSE298573, and the reviewer token is kzqvgquwhfatjcp. This manuscript does not report any original code. Any additional information required to reanalyse the data reported in this manuscript is available from the lead contact upon request.

## Results

### CMPortal analysis of metabolic variables in hiPSC-CM ischaemia modelling

To identify experimental variables associated with cardiomyocyte maturation and disease modelling, we analysed 322 hiPSC-CM differentiation protocols curated in the CMPortal database^27^. CMPortal applies decision tree–based feature importance analysis using Shannon entropy (information gain) and Gini impurity metrics to quantify associations between protocol variables and reported maturation or disease–modelling outcomes across curated hiPSC-CM studies. This analysis identified metabolic substrates strongly associated with maturation outcomes but markedly underrepresented in cardiovascular disease modelling studies (**Supplementary Figure 1a**). Among the 322 hiPSC-CM studies^27,42^, there are only a small number (29 studies) using metabolic manipulations, and few (9 studies) of these link to *in vitro* ischaemic modelling (**Supplementary Figure 1b**). These ischaemic modelling protocols employ a defined set of maturation strategies, highlighting opportunities to further define the role of protocol variables in optimising conditions for modelling cardiac ischaemia (**Table 1**). Further, CMPortal database shows that protocol variables associated with cardiomyocyte maturation showed limited overlap with those enriched in studies modelling ischaemic injury (**Supplementary Figure 2a**). Variables linked to improved contractile performance were primarily associated with extracellular matrix components and culture platform features, including gelatin or Matrigel coatings, ascorbic acid supplementation, Wnt inhibition, and three-dimensional culture systems, which are known to influence structural organisation and biomechanical maturation (**Supplementary Figure 2a, upper**). In contrast, protocols used for *in vitro* ischaemia modelling were enriched for metabolic media components such as KOSR, albumax, biotin, and carbohydrate substrates, together with defined culture media and prolonged Wnt inhibition (**Supplementary Figure 2a, lower**). These patterns suggest that while maturation studies commonly emphasise structural and developmental cues, disease-modelling protocols more frequently incorporate variables that influence cellular metabolic state.

**Table 1.** Characterisation of the ischaemic modelling protocols using hiPSC-CM in CMPortal^27^ database^*^.

| Reference | Differentiation protocol variables | hiPSC-CM backbone medium | Maturation strategies for hiPSC-CM | Hypoxia (or O<sub>2</sub> level) | pH (extracellular) | Duration (ISC+ REP) | LDH or cell stress measurement |
| --- | --- | --- | --- | --- | --- | --- | --- |
| Chen et al., 2019<sup>51</sup> | Cell line: WTC11<br>Matrix coating: Matrigel<br>hiPSC backbone: mTeSR-1<br>Seeding confluency (%): 92.5<br>Wnt activator and inhibitor: CHIR99021, Wnt-C59<br>Differentiation purity (%): n/a | RPMI 1640+2% B27<br>plus insulin | Bioreactor culture and extended culture | Anoxic gas (95% N<sub>2</sub>, 5% CO<sub>2</sub>) | ISC 6.4, REP 7.4 | ISC 6 h, REP 3 h | • LDH release<br>• Adenylate kinase<br>• Viability<br>• MMP staining<br>• ROS levels<br>• Intracellular pH |
| Sebastião et al., 2020<sup>9</sup> | Cell line: DF19-9-11T.H<br>Matrix coating: Matrigel<br>hiPSC backbone: mTeSR-1<br>Seeding confluency (%): 85<br>Wnt activator and inhibitor: CHIR99021 and Activin A, IWR1<br>Differentiation purity (%): 90 | RPMI 1640+2% B27 without insulin | T3 hormone-rich maturation medium in 3D aggregate culture | <0.4% in O<sub>2</sub> | ISC 6.8, REP 7.4 | ISC 5 h, REP 16 h | • Aggregates viability and diameter<br>• Molecular markers and ultrastructure<br>• Secretion of cytokines and growth factors<br>• Angiogenic potential |
| Hidalgo et al., 2018<sup>8</sup> | Cell line: C32<br>Matrix coating: Matrigel<br>hiPSC backbone: mTeSR<br>Seeding confluency (%): 100<br>Wnt activator and inhibitor: CHIR99021, IWP4<br>Differentiation purity (%): n.a | RPMI 1640+2% B27 plus insulin | DMEM, no glucose with 2% FCS, galactose, palmitic acid, oleic acid | 0% O<sub>2</sub> | ISC 6.2, REP 7.4 | ISC 2 h, REP 4 h | • LDH release<br>• PI/Hoechst staining<br>• ROS and caspase-3 staining |
| Friedman et al., 2024<sup>28</sup> | Cell line: WTC CRISPRi GCaMP<br>Matrix coating: Vitronectin XF<br>hiPSC backbone: mTeSR plus<br>Seeding confluency (%): 80<br>Wnt activator and inhibitor: CHIR99021, XAV-939<br>Differentiation purity (%): >80 | RPMI 1640+2% B27 plus insulin | Extended culture | 0.5% O<sub>2</sub> | ISC 7.4, REP 7.4 | ISC 18 h, REP 1 and 4 h | LDH release |
| Acun et al., 2019<sup>52</sup> | Cell line: HVRDi007-A, Huv-iPS4F1<br>Matrix coating: Geltrex<br>hiPSC backbone: mTeSR-1<br>Seeding confluency (%): n.a<br>Wnt activator and inhibitor: CHIR99021, IWP4<br>Differentiation purity (%): 93 | RPMI 1640+2% B27 plus insulin | Extended culture | 1% O<sub>2</sub> | ISC 7.4, REP 7.4 | ISC 48 h+REP 24 h; ISC 48 h+REP 2 h | Live/Dead fluorescence assay<br>ROS levels |
| Ellis et al., 2022<sup>53</sup> | Cell line: HVRDi007-A<br>Matrix coating: Geltrex<br>hiPSC backbone: mTeSR-1<br>Seeding confluency (%): 95<br>Wnt activator and inhibitor: CHIR99021, IWP4<br>Differentiation purity (%): n.a | RPMI 1640+2% B27 plus insulin | Extended culture | 0.1% O<sub>2</sub> | ISC 7.4, REP 7.4 | ISC 3 h, REP 3 h | Live/Dead assay<br>Cell length |
| Gao et al., 2018<sup>54</sup> | Cell line: in-house<br>Matrix coating: Matrigel<br>hiPSC backbone: n.a<br>Seeding confluency (%): n.a<br>Wnt activator and inhibitor: CHIR99021 | Glucose-depleted culture medium with abundant lactate | 3D culture with ECs and SCs | 1% O<sub>2</sub>, 5% CO<sub>2</sub>, 94% N<sub>2</sub> | ISC 7.4 | ISC 48 h | LDH release<br>TUNEL assay<br>Cell viability |
|  | Differentiation purity (%): 96 |  |  |  |  |  |  |
| Richards et al., 2020<sup>55</sup> | Cell line: iCell<br>Matrix coating: n.a<br>hiPSC backbone: n.a<br>Seeding confluency (%): n.a<br>Wnt activator and inhibitor: n.a<br>Differentiation purity (%): >95 | iCell Cardiomyocytes maintenance medium | Culture with FBs and HUVECs | 10% O<sub>2</sub> (organoid infarction model) | ISC 7.4 | ISC 10 days | • TUNEL assay |
| Peters et al., 2022<sup>7</sup> | Cell line: SCVI-273, UKKi032-C<br>Matrix coating: Matrigel<br>hiPSC backbone: E8<br>Seeding confluency (%): 85<br>Wnt activator and inhibitor: CHIR99021, Wnt-C59<br>Differentiation purity (%): 94 | RPMI 1640+2% B27 plus insulin | DMEM no glucose, low glucose, and fatty acids | 1% O<sub>2</sub> or 5% O<sub>2</sub> | ISC 7.4 | ISC 24 h or 4 h | • LDH release<br>• Cell viability<br>• TUNEL assay |

\*ISC, ischaemia; REP, reperfusion; LDH, lactate dehydrogenase; MMP, mitochondrial membrane permeability; ROS, reactive oxygen species; MI, myocardial infarction; DMEM, Dulbecco's modified eagle media; T3, triiodothyronine; FBS, fetal bovine serum; FCS, fetal calf serum; NCX1, sodium-calcium exchanger 1; FBs, human cardiac ventricular fibroblasts; HUVECs, human umbilical vein endothelial cells; CFs, cardiac fibroblasts; ECs, endothelial cells; SCs, smooth muscle cells.

To further assess the role of metabolic interventions, we examined associations between commonly used metabolic components and a range of structural, functional, and electrophysiological maturation metrics reported across the literature (**Supplementary Figure 2b**). Overall, metabolic variables showed limited and heterogeneous associations with these endpoints. In rare instances where we observed enrichment, it was most evident for fatty acid–based interventions, including palmitic acid and broader fatty acid or lipid supplementation. In contrast, widely used maturation cues such as T3 and IGF-1 showed minimal enrichment across the measured parameters (**Supplementary Figure 2b**), despite their widespread use in hiPSC-CM maturation protocols, suggesting that their contribution to functional maturation may be more context-dependent than previously assumed.

These observations motivated a systematic experimental assessment of the protocol variables commonly used when transitioning differentiated hiPSC-CMs into endpoint assays for ischaemia modelling. We therefore evaluated how replating conditions, culture substrates, backbone media, signalling modulators and metabolic supplementation (**Table 2**) influence cardiomyocyte maturation and susceptibility to ischaemic injury.

**Table 2.** A summary of experimental perturbations on hiPSC-CM maturation.

| Medium name | Base | Supplements |
| --- | --- | --- |
| RPMI+B27 | RPMI-1640 | 2% B27 with insulin |
| DMEM+FA | DMEM no glucose + 10 mM HEPES | 2 mM L-carnitine, 5 mM creatine, 5 mM taurine, 1 mM nonessential amino acids, 1 × insulin-transferrinselenium, 1 × linoleic-oleic acid. |
| DMEM+FBS | DMEM/F12 + 2% FBS | 2 mM L-glutamine, 0.1 mM nonessential amino acids, 0.1 mM β-mercaptoethanol, 50 u/mL penicillin/streptomycin. |
| StemPro-34 | StemPro-34 + 2 mM L-glutamine | 1 × nutrient supplement, 50 µg/mL ascorbic acid. |
| **Coating matrix** |  | Function |
| Vitronectin XF (VTN) |  | A recombinant human xeno-free extracellular matrix (ECM) protein. |
| Matrigel (MTG) |  | A mixture of ECM components such as laminin, type IV collagen and fibronectin. |
| **Signalling molecule** |  | Function |
| T3 |  | Thyroid hormone stimulation |
| Chetomin (CTM) |  | HIF-1a inhibition |
| Endothelin-1 (ET-1) |  | Hypertrophic remodelling promotion |
| Insulin-like growth factor 1 (IGF-1) |  | Growth hormone stimulation |
| Torin 1 (TO1) |  | mTOR signalling inhibitor |
| GW7647 (GW) |  | Fatty acid metabolism and β-oxidation activation |
| **Metabolic substrate** |  | Function |
| Palmitic acid (PA) |  | A saturated long-chain fatty acid |
| Linoleic-oleic acid (LOA) |  | Unsaturated fatty acids |
| PA+GW |  | A saturated long-chain fatty acid + uptake stimulation |
| LOA+GW |  | Unsaturated fatty acids + uptake stimulation |
| PA+2-Deoxy-D-glucose (2DG) |  | A saturated long-chain fatty acid + glycolysis inhibition |
| LOA+2DG |  | Unsaturated fatty acids + glycolysis inhibition |
| Galactose (Gal) |  | A monosaccharide sugar (glucose substitute) |

### Replating promotes cardiomyocyte maturation and increases susceptibility to ischaemia–reperfusion injury

We first examined whether replating hiPSC-derived cardiomyocytes influences physiological maturation and susceptibility to ischaemic injury. Cardiomyocytes generated using a high-density monolayer small-molecule differentiation protocol^25,26^ were replated and analysed over time in endpoint assays. Gene program analysis using scDRS^33^ showed that the baseline high-density monolayer differentiation protocol primarily enriched gene sets associated with cardiac function but showed limited enrichment for disease-associated programs linked to cardiometabolic disorders, including myocardial infarction and diabetes (**Figure 1a**).

Functional analysis using image-based contractility measurements (MUSCLEMOTION)^41^ demonstrated progressive improvement in contractile parameters following replating, including increased contraction amplitude and changes in peak-to-peak and time-to-peak kinetics (**Figure 1b** and **Supplementary Figure 3a**), consistent with enhanced physiological maturation.

To determine whether these maturation changes influence susceptibility to cellular stress, we evaluated replating density as a variable affecting ischaemia-induced injury using an *in vitro* ischaemia–reperfusion (IR) model^3^. Cardiomyocytes were exposed to hypoxia (0.5% O_, 18 hours) followed by normoxic reperfusion. Cells replated at lower seeding density in RPMI+B27 showed significantly increased LDH release following IR, indicating greater susceptibility to ischaemic injury (**Figure 1c**). In contrast, altering extracellular matrix substrate (Vitronectin XF versus Matrigel) did not significantly affect contractility (**Supplementary Figure 3b-d**) or IR-induced cell death under these conditions (**Supplementary Figure 3e**).

To define molecular changes associated with replating, we performed CAGE sequencing to identify differentially regulated transcriptional programs. Replating induced gene programs associated with muscle contraction and cardiac morphogenesis (**Figure 1d**). Compared with non-replated cells, replated cardiomyocytes showed increased expression of mature myofilament genes (*MYH7, MYL2, TNNI3*), hypertrophic growth regulators (*NPPB, IGF1, NPPA, MYOCD*), and electromechanical maturation genes (*SCN5A, CASQ2, CACNA2D1, KCNH2*), accompanied by reduced expression of glycolysis-associated genes (*HK2, GPI, GAPDH, ENO1*) (**Figure 1e**).

To examine the relevance of these transcriptional changes to human cardiac traits, we performed partitioned SNP heritability analysis using SBayesS^34^. Genes upregulated following replating were enriched for heritability associated with cardiac structure and function traits (RV and LV ejection fraction) but showed minimal association with loci linked to ischaemic heart disease or acute myocardial infarction (**Figure 1f** and **Supplementary Table 1**). Together, these results demonstrate that replating, particularly at lower seeding density, promotes molecular and functional maturation of hiPSC-derived cardiomyocytes and increases susceptibility to IR injury.

### Backbone medium determines metabolic maturation and susceptibility to ischaemia–reperfusion injury

We next examined whether the metabolic composition of the backbone culture medium influences cardiomyocyte maturation and susceptibility to ischaemic injury. Based on the CMPortal analysis, we compared two commonly used culture conditions: glucose-based RPMI+B27 and a fatty acid–containing medium lacking glucose (DMEM+FA) (**Supplementary Figure 1b, Figure 2a** and **Table 2**).

**Fig. 2.**
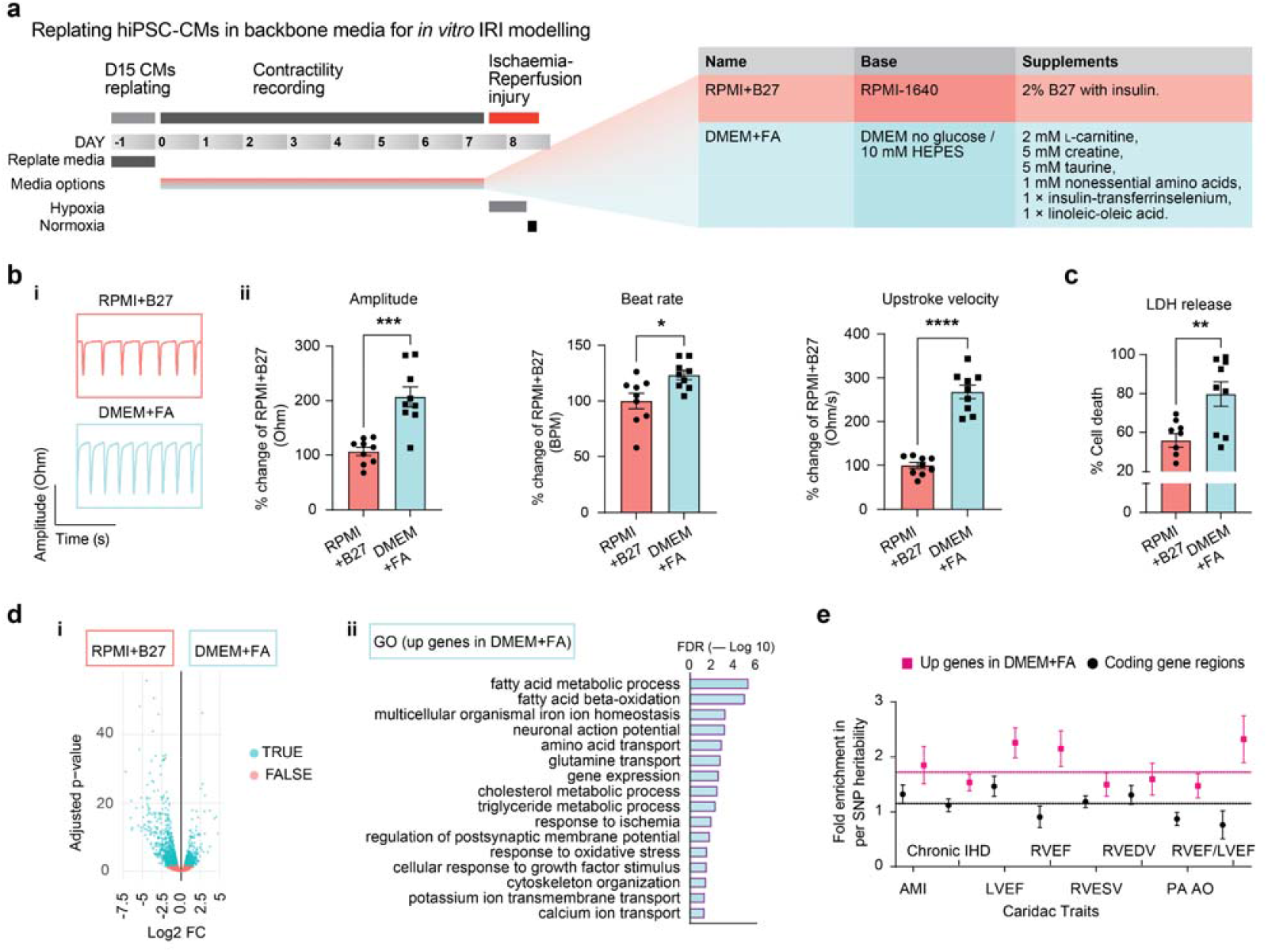
Comparative analysis of backbone medium influencing iPSC-CM maturation and ischaemic modelling. **(a)** Schematic of backbone medium by RPMI+B27 and DMEM+FA with supplements on maturity of hiPSC-CMs and IRI-induced stress responses. **(b)** An impedance-based contractility analysis using CardioExcyte 96 platform shows CM contractile performance in backbone medium RPMI+B27 or DMEM+FA with representative traces **(bi)**, and normalised peak values **(bii)** in amplitude, beat rate and upstroke velocity. **(c)** IRI-induced cell death comparing two backbone media conditioned CMs. **(d)** Molecular characterisation by RNA-sequencing on backbone medium comparison shows a volcano analysis of upregulated and downregulated genes in DMEM+FA compared to RPMI+B27 **(di)** and GO analysis of upregulated genes in DMEM+FA cultured CMs compared to RPMI+B27 conditions **(dii). (e)** Fold enrichment in per-SNP heritability analysis for up-regulated genomic regions in DMEM+FA-cultured than RPMI+B27-conditioned CMs versus all protein-coding genomic regions. AMI, acute myocardial infarction; Chronic IHD, chronic ischaemic heart disease; LVEF, left ventricular ejection fraction; RVEF, right ventricular ejection fraction; RVESV, right ventricular end-systolic volume; RVEDV, right ventricular end-diastolic volume; PA AO, pulmonary artery and aorta diameter ratio; RVEF/LVEF ratio, inverse-normalised right ventricular ejection fraction to left ventricular ejection fraction ratio. Unless otherwise specified, *n* = 3; 3 biological replicates, each with 3 technical replicates. Data are presented as mean ± SEM. Statistical analysis by two-tailed student’s *t*-test **(b-c)**. \**p*<0.05, \*\**p*<0.01, \*\*\**p*<0.001, \*\*\*\**p*<0.0001.

Compared with RPMI+B27, hiPSC-CMs cultured in DMEM+FA exhibited increased contraction amplitude, beat rate, and contraction velocity over a 7-day recording period (**Figure 2bi and 2bii**), indicating enhanced functional maturation. Among additional media tested, StemPro-34 also increased contractile amplitude and upstroke velocity relative to RPMI+B27, whereas DMEM/F12/FBS did not produce significant improvements (**Supplementary Figure 3f-h**).

We next examined whether these differences in maturation state altered susceptibility to ischaemia–reperfusion injury. Cardiomyocytes cultured in DMEM+FA showed significantly increased LDH release following IR injury compared with RPMI+B27 (**Figure 2c**). In contrast, StemPro-34 and DMEM/F12/FBS did not significantly alter IR-induced cell death relative to RPMI+B27 (**Supplementary Figure 3i**), indicating that enhanced contractility alone does not necessarily correspond to increased sensitivity to hypoxic injury.

Transcriptomic analysis of these different culture conditions revealed molecular programs associated with phenotypic differences. RNA-sequencing identified genes upregulated in DMEM+FA conditions that were enriched for fatty acid β-oxidation and pathways associated with cellular responses to ischaemia (**Figure 2di-dii**). Partitioned SNP heritability analysis using SBayesRC further showed that genes induced in DMEM+FA conditions were enriched for loci associated with ischaemic heart disease and acute myocardial infarction (**Figure 2e**), whereas genes upregulated in RPMI+B27 conditions were primarily associated with cardiac structure and function traits (**Figure 1f**).

Together, these results indicate that metabolic composition of the backbone culture medium shapes cardiomyocyte maturation state and transcriptional programs linked to ischaemia sensitivity. In particular, fatty acid–containing culture conditions promote a cardiomyocyte phenotype that exhibits both enhanced functional maturation and increased susceptibility to IR injury.

### Metabolic supplements enhance cardiomyocyte maturation and increase susceptibility to ischaemia– reperfusion injury in glucose-based media

We next evaluated whether metabolic and signalling factors commonly used to promote cardiomyocyte maturation influence susceptibility to ischaemic injury. **Figure 3a** summarises enrichment of protocol variables associated with maturation-related functional endpoints across studies using RPMI+B27. Matrix components such as Matrigel and extracellular matrix factors showed strong associations with structural and contractile maturation indicators, consistent with their established roles in cardiomyocyte organisation. Notably, several metabolic variables including fatty acids, palmitic acid, galactose, and related lipid and carbohydrate substrates were also enriched across contractile and calcium handling endpoints, suggesting that metabolic substrates represent recurrent features of protocols associated with improved cardiomyocyte maturation when cultured in RPMI+B27. These observations provided a rationale to systematically evaluate how defined metabolic and signalling cues influence cardiomyocyte maturation and susceptibility to ischaemic injury.

**Fig. 3.**
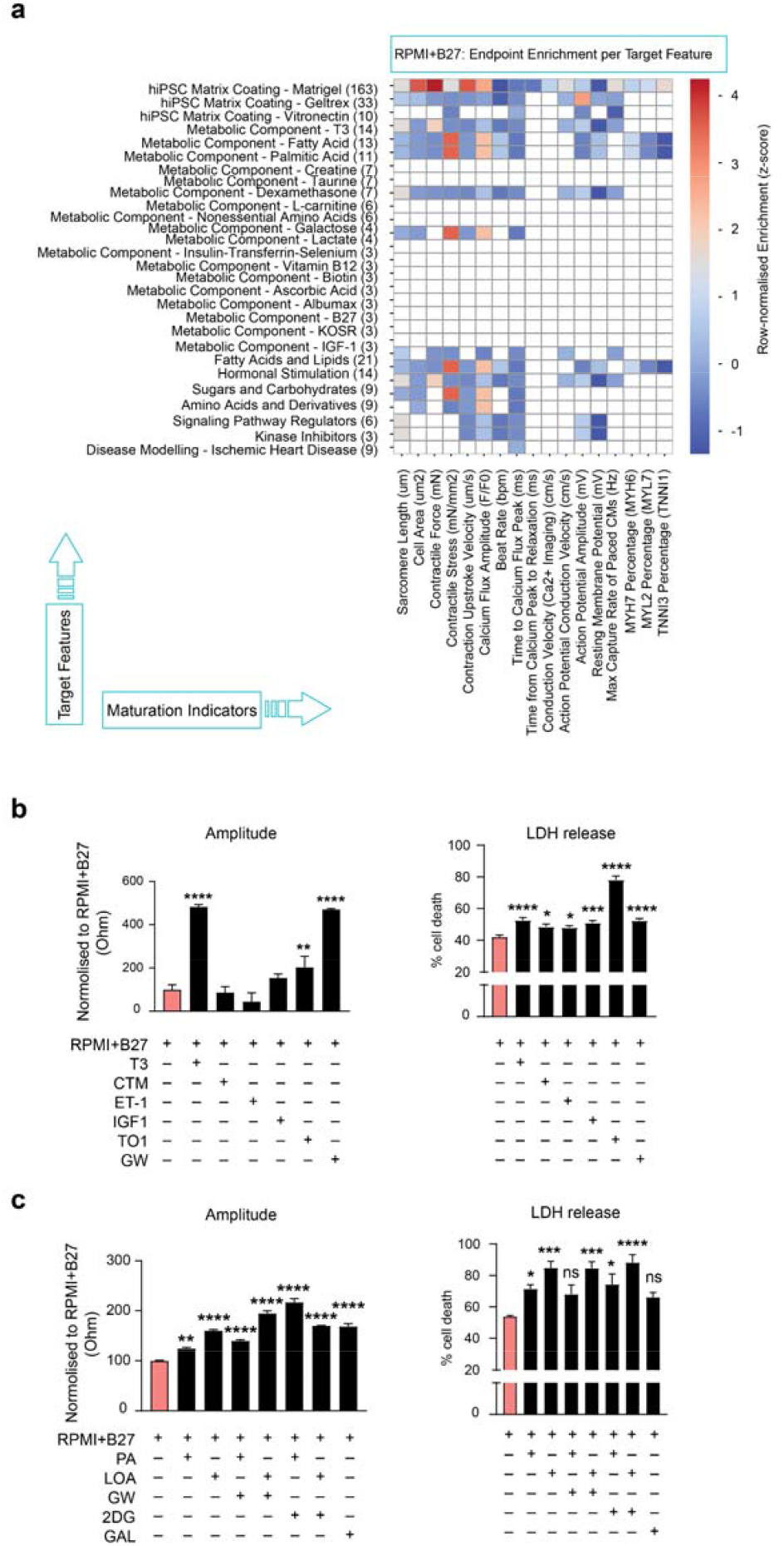
Evaluation of the interaction between RPMI/B27 medium and factors guiding modelling of ischaemic response. **(a)** Heatmaps illustrate the standardised z-scores for best available maturation values (columns) across protocol variables (rows) for studies using RPMI-1640+B27. **(b-c)** Fold change in peak amplitude of CMs cultured in RPMI+B27 and incubated with variables including signalling molecules **(b)** or metabolic substrates **(c)**, with the retrospective analysis of CMs susceptibility to cell death under ischaemia-reperfusion stress evaluating CMs cultured in RPMI+B27 and supplemented with signalling molecules **(b)** or metabolic substrates **(c)**. For **a**, colour represents standardised performance relative to within-media distributions: red indicates above-average maturation (positive z-scores), blue indicates below-average maturation (negative z-scores). Unless otherwise specified, *n* = 3; 3 biological replicates, each with 3 technical replicates. Data are presented as mean ± SEM. Statistical analysis by one-way ANOVA **(b, c)**. \**p*<0.05, \*\**p*<0.01, \*\*\**p*<0.001, \*\*\*\**p*<0.0001.

Guided by these enrichment patterns, we performed a targeted perturbation screen in which replated hiPSC-CMs were cultured in RPMI+B27 supplemented with individual maturation factors (**Table 2**). Contractile function was assessed using impedance-based recordings on the CardioExcyte platform^43-45^. Supplementation of RPMI+B27 with metabolic substrates (palmitic acid, linoleic–oleic acid, galactose) and signalling molecules (T3, GW7647, Torin1) significantly increased contraction amplitude, velocity, and beat rate compared with RPMI+B27 alone (**Figure 3b-c** and **Supplementary Figure 4a-b**).

We next tested whether these maturation changes influenced susceptibility to ischaemia–reperfusion injury. In most conditions, factors that enhanced physiological maturation also increased cell death following IR injury (**Figure 3b-c**). Together, these results indicate that metabolic and signalling cues used to promote cardiomyocyte maturation can also shift the metabolic state of hiPSC-CMs in ways that increase vulnerability to hypoxic stress.

### Metabolic backbone media determine the effectiveness of maturation supplements

Given that fatty acid–containing DMEM+FA enhanced cardiomyocyte maturation and increased susceptibility to ischaemia–reperfusion injury (**Figure 2)**, we next examined whether the maturation factors identified in RPMI+B27 conditions (**Figure 3)** further enhance maturation when applied in this metabolically defined medium.

Feature enrichment analysis from CMPortal suggested that, in contrast to RPMI+B27, relatively few protocol variables were associated with improved maturation endpoints when DMEM+FA was used as the backbone medium (**Figure 4a**). Functional validation confirmed this observation. Compared with RPMI+B27, DMEM+FA alone markedly increased cardiomyocyte contractility, whereas the addition of maturation supplements produced minimal further improvements in contractile performance (**Figure 4b** and **Supplementary Figure 4c**).

**Fig. 4.**
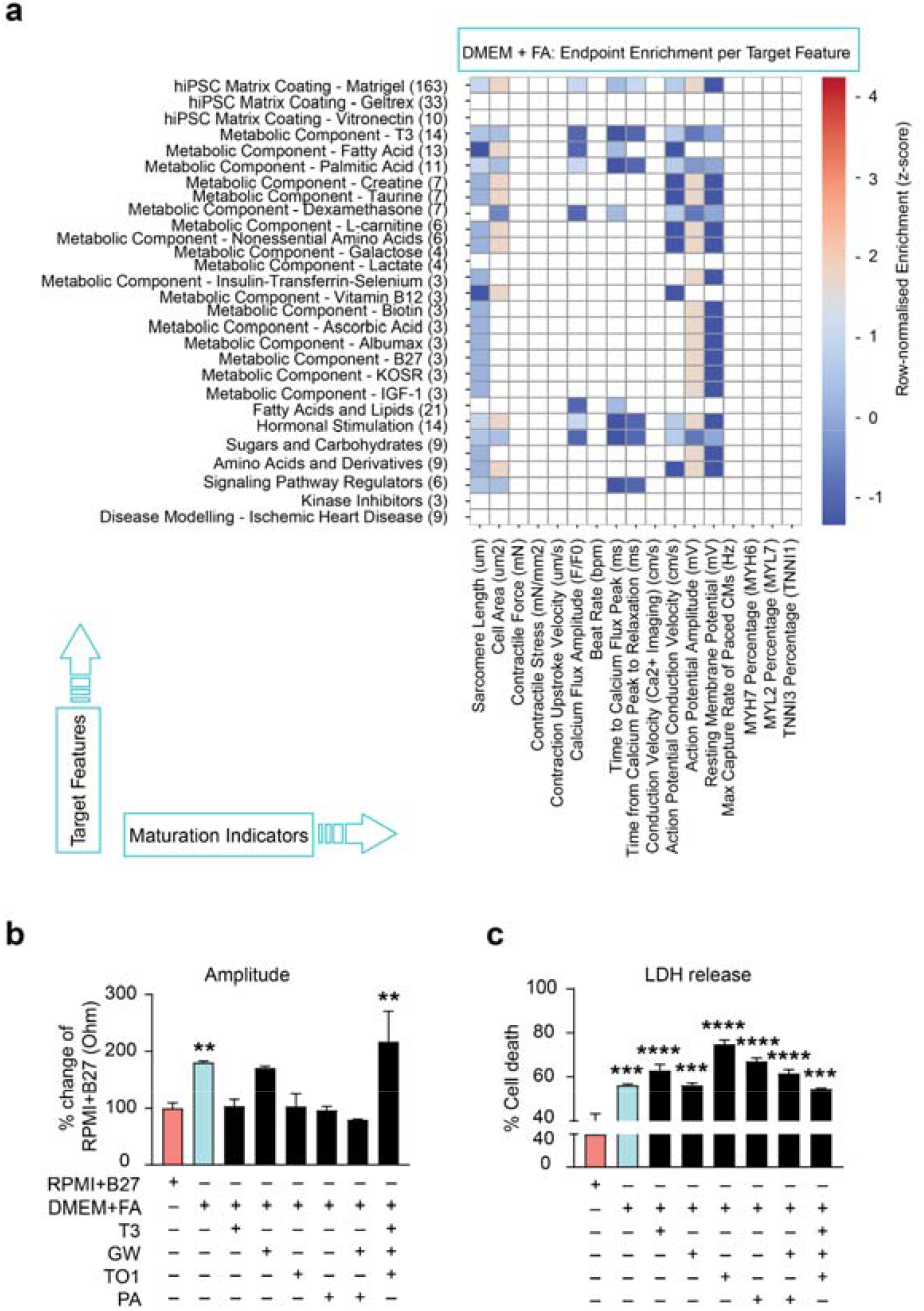
Evaluation of the interaction between DMEM/FA medium and factors guiding modelling of ischaemic response. **(a)** Heatmaps illustrate the standardised z-scores for best available maturation values (columns) across protocol variables (rows) for studies using DMEM+FA. **(b-c)** Fold change in peak amplitude **(b)** of CMs cultured in backbone medium RPMI+B27 or DMEM+FA and incubated with maturation factors, associated with analysis of CMs susceptibility to cell death **(c)** under ischaemia-reperfusion stress evaluating CMs cultured in backbone medium RPMI+B27 or DMEM+FA and incubated with maturation factors. For **a**, colour represents standardised performance relative to within-media distributions: red indicates above-average maturation (positive z-scores), blue indicates below-average maturation (negative z-scores). Unless otherwise specified, *n* = 3; 3 biological replicates, each with 3 technical replicates. Data are presented as mean ± SEM. Statistical analysis by one-way ANOVA **(b, c)**. \**p*<0.05, \*\**p*<0.01, \*\*\**p*<0.001, \*\*\*\**p*<0.0001.

We next examined whether these supplements altered susceptibility to ischaemia–reperfusion injury under DMEM+FA conditions. Consistent with the contractility results, most supplements did not further significantly increase IR-induced cell death beyond the level observed with DMEM+FA alone (**Figure 4c**).

Together, these findings demonstrate that the metabolic composition of the backbone culture medium establishes the baseline maturation state of hiPSC-derived cardiomyocytes and determines their responsiveness to additional maturation stimuli. In fatty acid–containing conditions, cardiomyocytes exhibit a metabolically mature phenotype that enhances functional performance and increases susceptibility to ischaemic injury, with minimal additional benefit from maturation supplements.

## Discussion

This study identifies metabolic environment as a central determinant of cardiomyocyte susceptibility to ischaemia– reperfusion injury in human stem cell–derived models. By integrating literature mining with systematic perturbation of experimental variables used during endpoint assay preparation, we show that protocol choices made after cardiomyocyte differentiation, including replating density, culture substrate, backbone medium, and metabolic supplementation, reshape cardiomyocyte maturation state and influence injury responses. In particular, culture conditions that promote metabolic maturation toward fatty acid utilisation enhance contractile performance while simultaneously increasing susceptibility to ischaemic injury. These findings establish a direct relationship between metabolic maturation and injury responsiveness in hiPSC-derived cardiomyocytes and highlight the importance of experimental design in shaping disease modelling outcomes.

This work demonstrates that cardiomyocyte state is strongly influenced by the transition from differentiation protocols to endpoint assay conditions used for disease modelling. Replating of hiPSC-CMs induced transcriptional and functional changes consistent with physiological maturation, including increased expression of mature sarcomeric and electrophysiological genes and reduced expression of glycolytic pathways. Importantly, these changes were accompanied by increased susceptibility to ischaemia–reperfusion injury, particularly at lower replating densities. These findings indicate that experimental technical variables can substantially reshape cardiomyocyte physiology and therefore influence the utility and interpretation of disease modelling studies. This observation highlights the need to consider the entire experimental workflow when designing human stem cell models of cardiac injury, and aligns with recent reports challenging the assumption that a single “optimal” maturation protocol universally enhances physiological relevance, instead emphasising that the effects of culture conditions on disease phenotypes are highly customised^27,42^.

Second, we show that metabolic composition of the culture environment acts as a dominant regulator of cardiomyocyte maturation and injury sensitivity. Fatty acid–containing DMEM+FA medium promoted both enhanced contractile maturation and increased susceptibility to ischaemia–reperfusion injury compared with glucose-based RPMI+B27 conditions. Transcriptomic analysis further revealed activation of fatty acid oxidation pathways and enrichment of genes associated with ischaemic heart disease risk loci. These results are consistent with the metabolic shift that occurs during postnatal cardiac maturation, in which cardiomyocytes transition from glycolysis toward mitochondrial fatty acid oxidation^46-48^. Our findings therefore demonstrate that metabolic substrate availability alone can drive a cardiomyocyte state that more closely reflects the energetic vulnerability of adult myocardium.

Lastly and most unexpectedly, we show that the metabolic backbone medium determines the effectiveness of additional maturation stimuli. While metabolic supplements and signalling factors improved contractile performance and increased injury sensitivity when applied in glucose-based RPMI+B27 conditions, these interventions produced minimal additional effects when cardiomyocytes were cultured in fatty acid–containing DMEM+FA medium. This finding suggests that the metabolic environment establishes a baseline maturation state that constrains the responsiveness of cardiomyocytes to further maturation cues. In other words, metabolic context operates hierarchically, with backbone media composition exerting greater influence on cardiomyocyte state than individual maturation supplements. This conceptual advance in a hierarchical metabolic context is supported by recent studies demonstrating that the metabolic substrate composition of culture media is a major driver of cardiomyocyte metabolic state and functional maturation^6,49,50^.

Together, these findings highlight the importance of metabolic context in determining the physiological relevance of hiPSC-derived cardiomyocyte models. For modelling ischaemia–reperfusion injury, metabolic substrate composition and cardiomyocyte energetic state appear to be key determinants of injury fidelity. By defining culture conditions that link metabolic maturation with physiologically relevant injury responses, this study provides a foundation for improving the reproducibility and translational utility of human stem cell models for studying cardiac ischaemia and for evaluating cardioprotective therapies.

## Acknowledgements

This work was supported by funding from the National Heart Foundation (106721 to NP) and the MRFF (MRFCDDM000033 to NP). We acknowledge Dr Meredith A. Redd for providing her suggestions on the experimental design of maturation strategies and sensitivity to cardiac ischaemia. We also acknowledge Dr Clayton E. Friedman for assisting with CAGE-sequencing experiments and data collection.

## Author contributions

YC conceptualised and designed the study, executed the experiments, performed data analysis, and drafted the manuscript, serving as co-corresponding author and intellectual lead for the project. CC contributed to the CMPortal database analysis and feature prediction, informing physiological maturation performance. SN contributed to scDRS and heritability analysis. WS assisted in processing bulk RNA-sequencing data and performing DE gene analysis for a backbone medium comparison. SS and Friedman contributed to CAGE-sequencing data generation and the volcano plot in replicating versus non-replicating cardiomyocytes. Fang contributed to cardiac differentiation from human pluripotent stem cells and quality control. NP contributed to the experimental design, data interpretation, manuscript preparation and overall project supervision.

