## Supplementary Information for "Hierarchical control of cardiomyocyte maturation and ischaemia sensitivity by metabolic culture conditions"

### Supplementary figures and legends

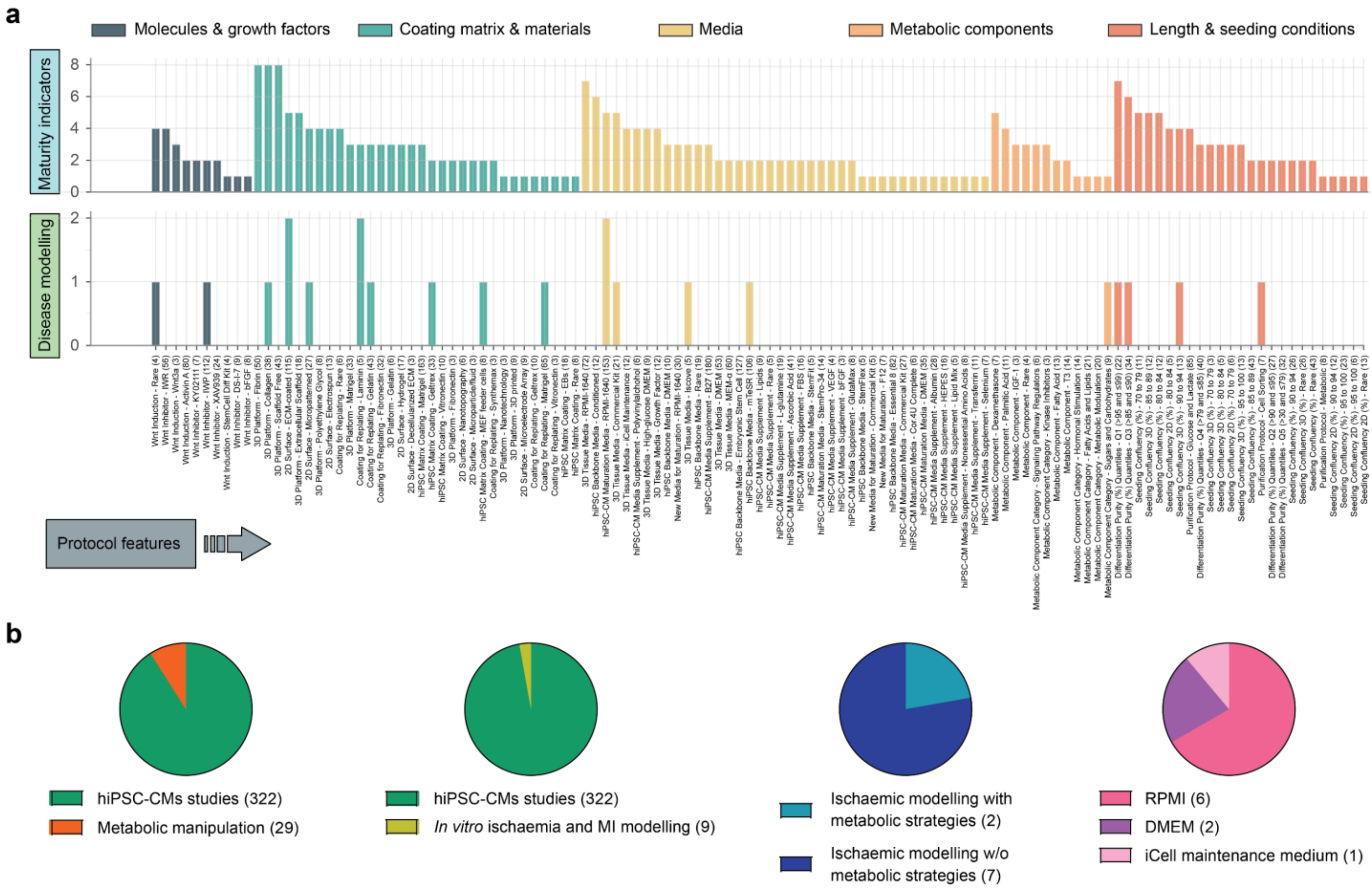

**Supplementary Figure 1 | Quantitative analysis of CMPortal database to evaluate protocol variables associated with physiological maturation and disease modelling. (a)** Bar plots showing 5 categories of protocol variables significantly associated with best maturity outcomes (upper) and modelling purposes for diseases (lower) of 322 protocols. Significance was defined as $p$-values < 0.01 using either the Gini or Entropy metrics in CMPortal permutation tests. Protocol variable categories include molecules and growth factors (dark green), coating matrix and material (green), selection of media (yellow), metabolic substrates (orange), and other considerations of protocol length and seeding conditions (red). Maturity outcomes span structural (sarcomere length, cell area), functional (contractile properties, calcium handling, electrophysiology), and molecular (gene and protein expression) maturation parameters. The Y-axis represents the number of protocols in which each protocol variable category (functional group) was significantly enriched for either improved maturity outcomes (upper plot) or specific disease modelling applications (lower plot), based on $p < 0.01$ significance; disease modelling refers to the specific cardiac disease categories tested in the protocols, including genetic disorders, arrhythmic conditions, ischaemic infarction, dilated and hypertrophic cardiomyopathy, and fibrotic diseases. Each bar represents the number of protocols in which a given functional group was significantly enriched for that disease application. **(b)** Pie charts show the number of studies using metabolic strategies, ischaemic modelling among the 322 hiPSC-CM database. In *in vitro* ischaemic modelling studies, pie charts show the metabolic versus other strategies, as well as the most common backbone media used in these protocols.

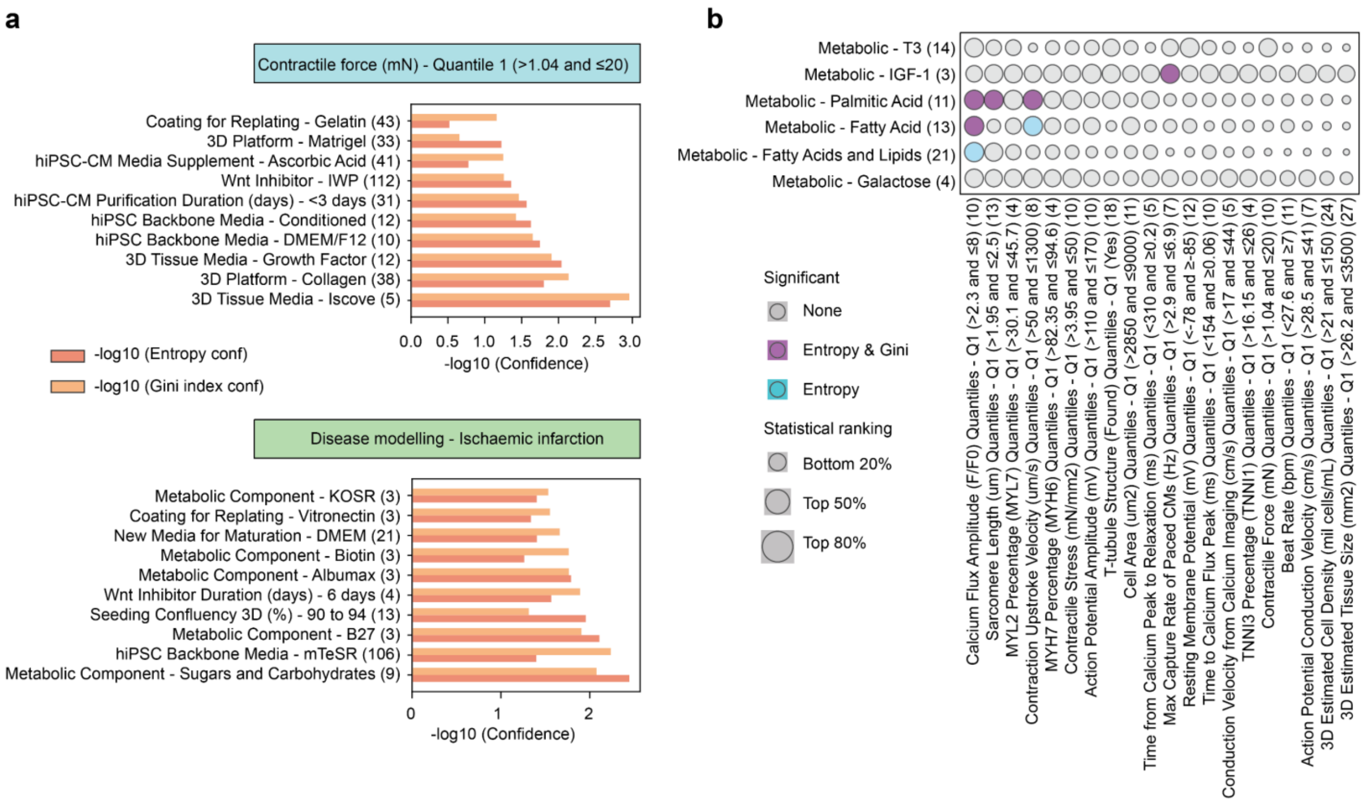

**Supplementary Figure 2 | Quantitative analysis of CMPortal database to evaluate protocol variables associated with physiological contractility and ischaemia modelling. (a)** Horizontal bar plots showing the top 10 enriched protocol variables for contractile strength (upper) and ischaemic modelling (lower). The bar height indicates the association significance based on entropy (red), and Gini index (orange). **(b)** Bubble plot analysis of common metabolic component rankings of their significance towards maturation indicators. Bubble size is proportional to inverse rank score (larger bubbles indicate higher-ranking associations). Colour coding represents statistical significance: gray (non-significant in either metric), blue (entropy-significant), purple (significant in both entropy and gini index). Statistical analysis by feature importance with Shannon Entropy (information gain via the reduction of entropy by splitting the dataset with the feature) and Gini Impurity (the random probability that the target functional outcome, is incorrectly miscategorised in a subset after splitting with the protocol category **(a**, **b)**.

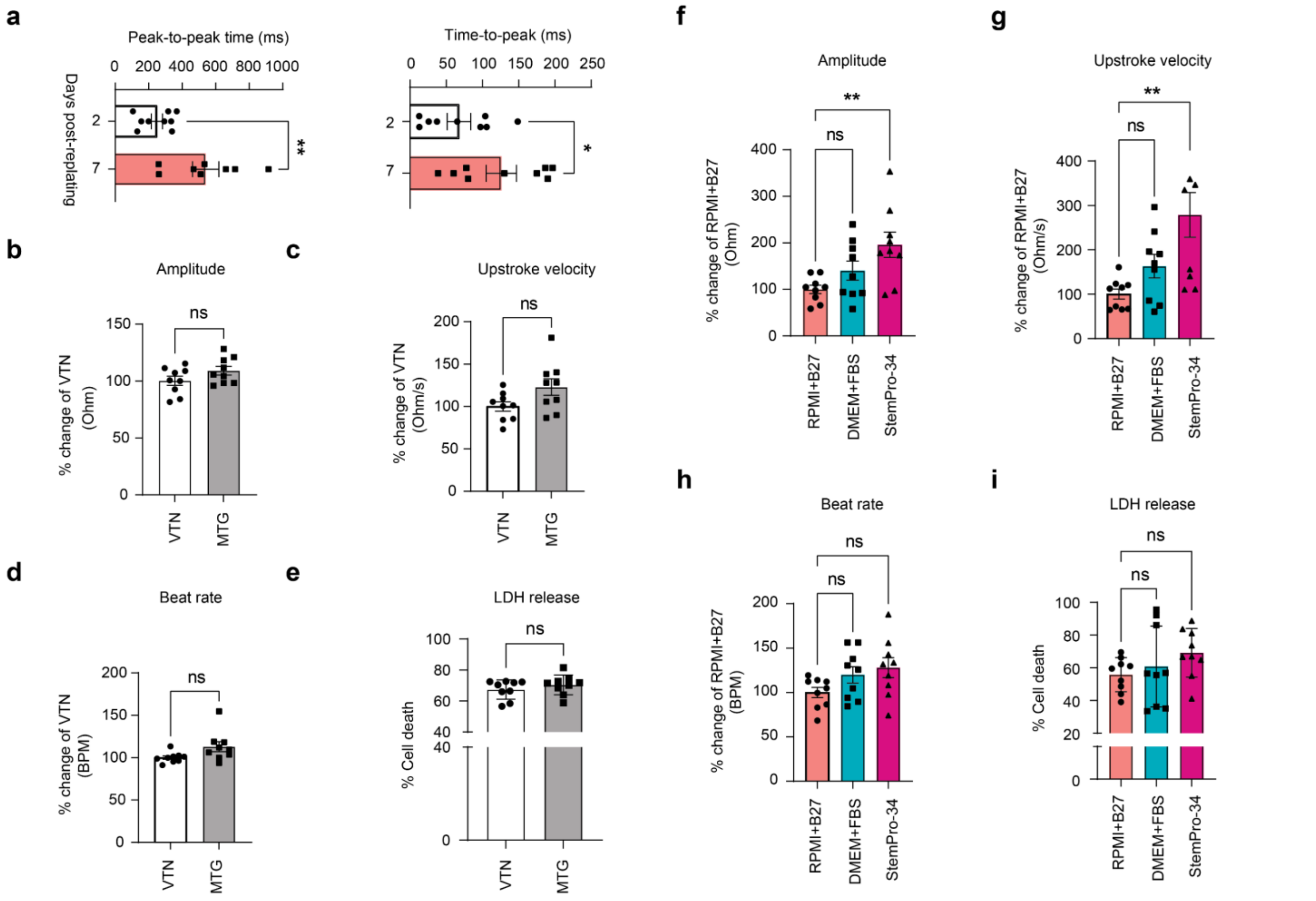

**Supplementary Figure 3 | Replating, matrix and media effects on hiPSC-CMs functional contractility and cardiac cell death. (a)** MUSCLEMOTION contractility analysis of cardiomyocytes post-replating 2 days and 7 days. **(b-e)** Analysis of maturity CMs on contractility **(b-d)** and cell death under ischaemia-reperfusion stress induced by coating matrix **(e)**. **(f-h)** Analysis of maturity CMs on amplitude **(f)**, upstroke velocity **(g)**, and beat rate **(h)** by comparing media options, including RPMI+B27, DMEM/F12/FBS, and StemPro-34. **(i)** LDH release assay to detect cell death under ischaemia-reperfusion stress, comparing hiPSC-CMs cultured in different media types. ($n$ = 3; 3 biological replicates, each with 3 technical replicates). Data are presented as mean ± SEM. Statistical analysis by two-tailed student's $t$-test **(a-e)**, or one-way ANOVA **(f-i)**. *$p$<0.05, **$p$<0.01, ***$p$<0.001, ****$p$<0.0001.

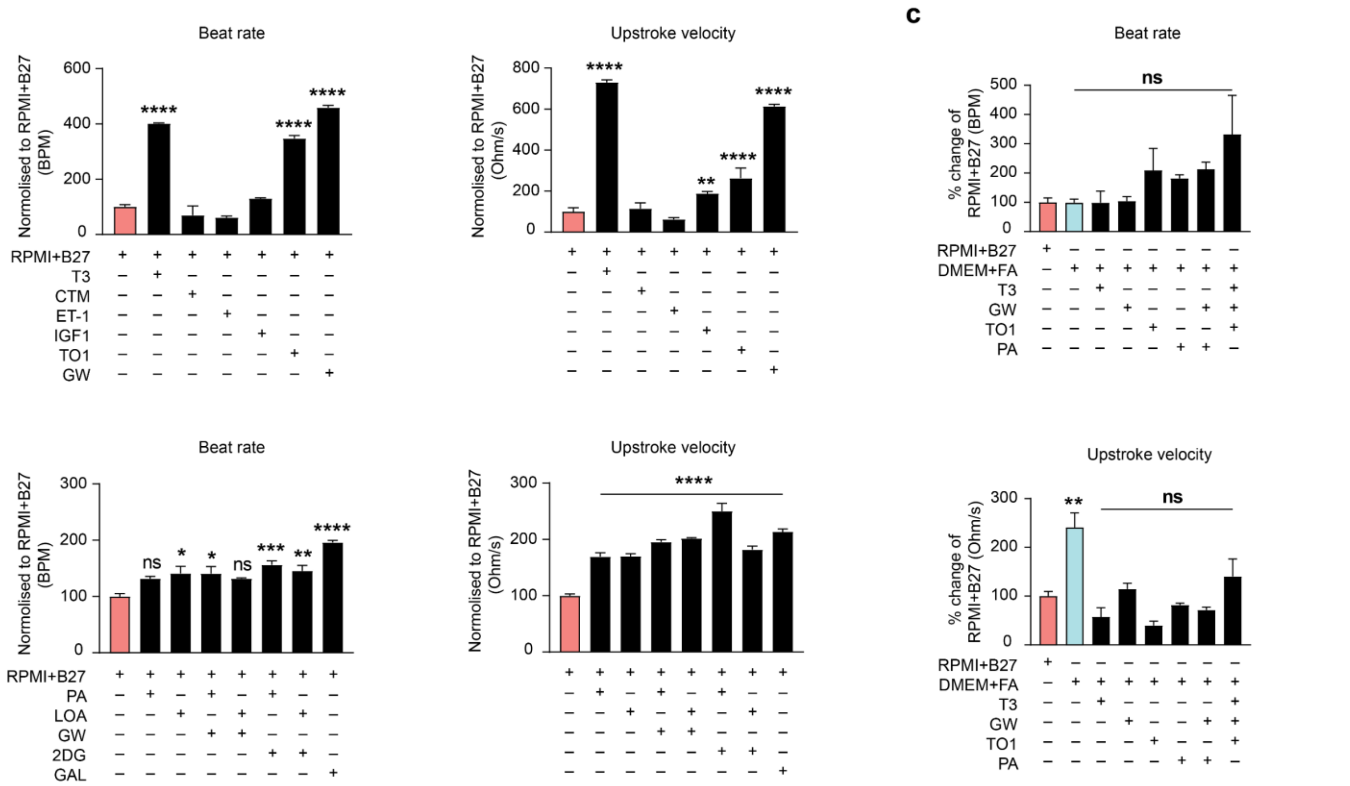

**Supplementary Figure 4 | Functional characterisation of hiPSC-CMs maturation by molecules, metabolic substrates and selective culture conditions. (a-b)** Analysed fold changes of beat rate and upstroke velocity of CMs cultured in backbone medium RPMI+B27 incubated with signalling molecules **(a)** and metabolic regulators **(b)**. **(c)** Analysed fold changes of beat rate, upstroke velocity of CMs cultured in backbone medium DMEM+FA supplemented with selective maturation factors ($n = 3$; 3 biological replicates, each with 3 technical replicates). Data are presented as mean ± SEM. Statistical analysis by one-way ANOVA **(a-c)**. *$p<0.05$, **$p<0.01$, ***$p<0.001$, ****$p<0.0001$.

**Reagent or resource table**

| Reagent or resource | Source | Identifier |
|---|---|---|
| Antibodies | | |
| Cardiac Troponin T | Thermo Fisher Scientific | Cat. #MA5-12960 |
| Phycoerythrin (PE)-conjugated sarcomeric α-actinin (SA) | Miltenyi Biotec Australia Pty | Cat. #130-123-773 |
| PE-conjugated anti-human isotype (IgG) | Miltenyi Biotec Australia Pty | Cat. #130-119-964 |
| Chemicals, peptides, and recombinant proteins | | |
| Vitronectin XF™ | STEMCELL Technologies | Cat. #07180 |
| mTeSR™ Plus maintenance medium | STEMCELL Technologies | Cat. #100-0276 |
| Rock inhibitor Y-27632 (ROCKi) | STEMCELL Technologies | Cat. #72308 |
| Bovine serum albumin (BSA) | Sigma-Aldrich | Cat. #A9418-50G |
| L-Ascorbic acid 2-phosphate sesquimagnesium salt hydrate (ascorbic acid, AA) | Sigma-Aldrich | Cat. #A8960-5G |
| CHIR-99021 (CHIR) | STEMCELL Technologies | Cat. #72054 |
| XAV-939 | STEMCELL Technologies | Cat. #72674 |
| RPMI-1640 medium | Thermo Fisher Scientific | Cat. #11875093 |
| B-27 supplement plus insulin (50X, B27/insulin) | Thermo Fisher Scientific | Cat. #17504001 |
| Fetal bovine serum (FBS) | ThermoFisher Scientific | Cat. #10099141 |
| 2.5% Trypsin | Thermo Fisher Scientific | Cat. #15090046 |
| Dimethyl sulfoxide | Sigma-Aldrich | Cat. #2650 |
| Saponin | Sigma-Aldrich | Cat. #S7900 |
| 4% paraformaldehyde | Sigma-Aldrich | Cat. #158127-5G |
| DMEM no glucose medium | Thermo Fisher Scientific | Cat. #11966025 |
| MEM Non-essential amino acids | Thermo Fisher Scientific | Cat. #11140050 |
| L-Carnitine Hydrochloride | Merck Life Science Pty Ltd | Cat. #C0283-5G |
| Insulin-Transferrin-Selenium (100X) | Thermo Fisher Scientific | Cat. #41400045 |
| Linoleic Acid-Oleic Acid-Albumin (100X) | Merck Life Science Pty Ltd | Cat. #L9655-5ML |
| Taurine | Sigma-Aldrich | Cat. #T8691-25G |
| Creatine, anhydrous | Sigma-Aldrich | Cat. #C0780-50G |
| Triton X-100 | Sigma-Aldrich | Cat. #X100 |
| Versene solution | Thermo Fisher Scientific | Cat. #15040066 |
| HEPES solution (10 mM) | Sigma-Aldrich | Cat. #H0887-100ML |
| Critical commercial assays | | |
| LDH cytotoxicity detection kit | Roche | Cat. #11644793001 |
| Deposited data | | |
| CAGE-sequencing data | Friedman et al[1] | ArrayExpress: E-MTAB-11719 |
| Bulk RNA-sequencing data | This manuscript | GEO: GSE298573 |
| hPSC-CM foundational database | Ewoldt et al[2] | DOI: https://doi.org/10.1038/s41592-024-02480-7 |
| CMPortal dataset | Chow et al[3] | https://palpantlab.com/cmportal |
| Experimental models: Cell lines | | |
| Human iPSC: WTC-11, IST3323, TOB421 | WTC-11 is a Gift from Gladstone Institute of Cardiovascular Disease, UCSF[4,5] | RRID: CVCL_Y803 |
| Software and algorithms | | |
| Python | Python Software Foundation | v3.12 |
| Pandas | pandas.pydata.org | v1.3.5 |
| NumPy | numpy.org | v1.21.0 |
| Flask | flask.palletsprojects.com | v2.0.1 |

| | | |
|---|---|---|
| Gunicorn | gunicorn.org | v20.1.0 |
| Nginx | nginx.org | v1.27.4 |
| htseq-count | HTSeq-count software | https://htseq.readthedocs.io/en/latest/ |
| STAR aligner | STAR | https://github.com/alexdobin/STAR |
| GraphPad Prism 9 (Version: 9.3.1) | GraphPad Software | https://www.graphpad.com/ |
| FACSDiva software | BD Biosciences | https://www.bdbiosciences.com/en-au/products/software/instrument-software/bd-facsdiva-software |
| FlowJo software (Version: 10.6.2) | Tree Star | https://www.flowjo.com/solutions/flowjo |
| DESeq2 | Love et al[6] | https://bioconductor.org/packages/release/bioc/html/DESeq2.html |
| CardioExcyte Control96 (Version: 1.4.7.1) | Nanion Technologies GmbH | https://www.nanion.de/products/cardioexcyte-96/ |
| Other | | |
| CardioExcyte 96 system | Nanion Technologies GmbH | https://www.nanion.de/products/cardioexcyte-96/ |
| CardioExcyte NSP-96 plates | Nanion Technologies GmbH | Cat. #201001 |
| FACS CANTO II system | Becton Dickinson | https://www.bdbiosciences.com/en-au/products/instruments/flow-cytometers/clinical-cell-analyzers/facscanto |
| LightSail | Amazon Web Services | https://aws.amazon.com/lightsail/ |
| 96-well culture plates | Nunc | Cat.#150318 |
| 0.22um filter | Sigma-Aldrich | Cat. # CLs431218-50EA |
| RNeasy Mini Kit | QIAGEN | Cat. #74106 |

**Resource availability**

Further information and requests for resources and reagents should be directed to and will be fulfilled by the Lead Contact, Nathan Palpant. This study did not generate new unique reagents.
